# Formation of hemiclonal reproduction and hybridogenesis in *Pelophylax water* frogs studied with species-specific cytogenomic probes

**DOI:** 10.1101/2023.10.29.564577

**Authors:** Choleva Lukáš, Doležálková-Kaštánková Marie, Labajová Veronika, Sember Alexandr, Altmanová Marie, Lukšíková Karolína, Chung Voleníková Anna, Dalíková Martina, Nguyen Petr, Pustovalova Eleonora, Fedorova Anna, Dmitrij Dedukh

## Abstract

Meiosis is a conservative process in all sexual organisms which ensures fertility and is central for producing genetic diversity by recombination and random segregation of parental chromosomes. Yet unexplored mechanisms may disrupt it and cause ‘loss of sex’ followed by the emergence of clonal modes of reproduction. Interspecific hybridization is the primary trigger for this process, but mechanistic basis of the transition to asexuality remains still unknown for most vertebrate animals. To study these processes in water frogs, we performed reciprocal mating between two sexual species, *Pelophylax ridibundus* and *P. lessonae*, and produced vital F1 progeny (*P. esculentus*). The RepeatExplorer2 analysis of low-coverage genomic data of the two parental species identified the *P. lessonae*-specific minisatellite marker *PlesSat01-48* (44 bp), which hybridized to (peri)centromeric regions of two chromosome pairs in *P. lessonae* – the acrocentric chromosome 8 and the chromosome 10 (a carrier of nucleolar organizer region; NOR). Chromosomal mapping combining the novel hybridization probe with the previously designed marker for *P. ridibundus*-specific centromeric satellite DNA showed that the *P. esculentus* progeny do not reproduce sexually. Instead, the F1 generation of *P. esculentus* instantly modified its gametogenesis and established asexual reproduction via hybridogenesis. Gametogenic modifications included premeiotic elimination of one of the parental genomes and clonal propagation of the remaining genome via endoreplication followed by standard meiotic division. The origin of DNA elimination and hybridogenesis in laboratory-produced hybrids supports a hypothesis that *P. esculentus* arises recurrently in nature whenever parental species come into reproductive contact. Based on the observed pattern of DNA elimination in the F1 progeny we discuss the origin and evolution of population systems in water frogs and the applicability of a newly designed chromosomal probe for other *Pelophylax* taxa.

## INTRODUCTION

Meiosis is a fundamental process in all sexual organisms as it ensures fertility and facilitates genetic diversity by recombination and random segregation of parental chromosomes (Grandont et al., 2013; Mercier et al., 2015; Zielinski, M. L., & Mittelsten Scheid, O., 2012). Although the meiotic division is a highly conserved process across eukaryotes (Bengtsson, 2009), it can be disrupted by various mechanisms, which can result in ‘loss of sex’ and the emergence of asexual modes of reproduction (Lamatsch & Stöck, 2009; Schurko et al., 2009). Interspecific hybridization is the primary trigger for this process, however, the mechanistic basis of the transition to asexuality is still poorly understood for most vertebrate animals (Neiman et al., 2014).

The association between hybridization and asexuality raised the question of whether the production of a clonal gamete by a diploid ancestor precedes or follows the initial hybridization event (Avise et al., 1992; Beukeboom & Vrijenhoek, 1998). The basis of the problem when studying this phenomenon in detail lies in the fact that most of the asexual taxa have originated from one or a few distinct hybridization events in the past (Neiman et al., 2014). The evidence suggesting that only a subset of possible parental genome combinations may induce asexuality came largely from studies conducted on laboratory-produced hybrid offspring of sexual animal species that are thought to be analogous to the sexual progenitors of extant asexual hybrids (Neiman et al., 2014). Unfortunately, these synthetic hybrids were mostly sexual or infertile (e.g. Cole et al., 2010; Stöck et al., 2010), and those being asexual were often characterized by low fertility and/or survivorship relative to naturally produced hybrid asexuals (Wetherington et al., 1987). Hybridogenetic water frogs of the genus *Pelophylax* represent a vertebrate tetrapod allowing for repeated laboratory synthesis of viable and fertile asexual hybrids (Berger, 1968; Hotz et al., 1985), providing thus an excellent opportunity to examine changes in mechanisms and regulation of meiotic division *in situ* during the switch from sexual to asexual reproductive mode.

The model system of hemiclonal water frogs includes *Pelophylax esculentus* (Linnaeus, 1758), the edible frog (of a typical genomic composition LR), which is a hybrid taxon originally emerging from matings between *Pelophylax lessonae* (Camerano, 1882), the pool frog (LL), and *Pelophylax ridibundus* (Pallas, 1771), the marsh frog (RR). Hybridogenetic *P. esculentus* must mate at each generation with a sexual host who provides the genome, which is being repeatedly excluded. In most of the species’ European distribution range, diploid hybrids live in sympatry with the parental species *P. lessonae*. In these so-called L-E systems, a hybrid *P. esculentus* excludes its haploid L genome, transmits only the haploid R genome in its gametes, and restores the hybrid genome composition in the new generation through mating with *P. lessonae*. In some populations, however, the process is reversed (the R-E system) as most *P. esculentus* hybrids exclude and later regain the R genome (by mating with *P. ridibundus*), while transmiting the L genome (reviewed by Graf et Polls Pelaz, 1989; Plötner, 2005). *Pelophylax esculentus* is further exceptional in the production of both fertile females and males in L-E populations, and very often, all-male lineages are present in the R-E system, producing only L or both L and R gametes (referred to as amphispermy; Uzzell et al., 1977; Vinogradov et al., 1991). Both systems have been found in Central Europe, typically as all-diploid populations (Hoffmann et al., 2015).

Hybridogenesis is distributed from angiosperms to invertebrates and vertebrate animals (Lavanchy & Schwander, 2019). It involves transmitting only one of the parental genomes to gametes, while the other is eliminated during gametogenesis and regained by mating with the complementary species. During the key process of programmed DNA elimination, the chromosomes of a particular parent are removed in a strictly selective manner, however, the exact timing of this process varies among taxa and takes place either during gonial cell divisions or during meiosis (Dedukh & Krasikova, 2022). In the field-caught water frogs, it happens premeiotically during the early gametogenesis (Chmielewska et al., 2018, 2022; Dedukh et al., 2017, 2019, 2020), and either via the formation of micronuclei around epigenetically tagged chromosomes which lag during mitotic anaphase or through a ‘chromosome budding’ out of the interphase nucleus (Chmielewska et al., 2018, 2022; Dedukh et al., 2019, 2020; Ogielska, 1994). So-far performed studies related to DNA elimination were marker-limited and consisted of monitoring the presence or absence of a *ridibundus*-specific chromosome marker which is entirely absent from *lessonae* chromosomes. This was achieved either by AMD/4′,6-diamidino-2-phenolindole (DAPI) staining (Bucci et al., 1990; Heppich et al., 1982) or chromosomal mapping of a pericentromeric tandemly repeated *RrS1* sequence (Ragghianti et al., 1995). Use of an interstitial telomeric site (ITS), which is present exclusively on the chromosome pair bearing nucleolus organizer regions (NOR), applies to metaphase stage only and requires high-quality chromosome spreads (Dedukh et al., 2013, 2015), while a genomic in situ hybridization (GISH) (Zaleśna et al., 2011) and comparative genome hybridization (CGH) (Doležálková et al., 2016), i.e., the application of whole-genomic probes, have their specific limitations in case of studying interphase cells and gametes. Therefore, searching for other reliable cytogenetic markers with a high predictive value and reproducibility is crucial to advance investigations of hybridogenesis and DNA elimination in *Pelophylax* water frogs.

Designing markers for closely related species often represents a challenge due to sequence homology. The implementation of bioinformatic pipelines allowing for de novo repeat identification in low-pass genome sequencing data greatly enhanced our ability to identify and classify previously hardly tractable repeats such as satellite DNA (satDNA) and mobile elements (e.g. Negm et al., 2021; Novák et al., 2020; Vondrak et al., 2020). Particularly satDNA is a tandemly-repeated class known to undergo rapid sequence evolution in a genome (Garrido-Ramos, 2017; Plohl et al., 2012; Thakur et al., 2021). As a consequence, closely related species and even conspecific populations might greatly differ in their satDNA landscapes (Ávila Robledillo et al., 2020; Bracewell et al., 2019; Feliciello et al., 2014) which can be leverated for the identification of parental genomes in hybrid/polyploid individuals (e.g., Belyayev et al., 2018; Heitkam et al., 2020; Marta et al., 2020; T. Schmidt et al., 2019) and in diverse evolutionary studies (e.g., da Silva et al., 2020; Gatto et al., 2018, 2021; Smith et al., 2020; Vittorazzi et al., 2014).

In the present study, we successfully designed a genomic marker for *P. lessonae* genome which is hence complementary to the already available *RrS1* marker for *P. ridibundus* provided by Ragghianti et al., (1995). Combining these two markers, we investigate parental genomes during gametogenesis in laboratory-produced hybrid *P. esculentus*, and de novo origin of its programmed DNA elimination and hybridogenesis.

## MATERIAL AND METHODS

### Sampling and taxa determination

*Pelophylax* water frog specimens belonging to seven diploid sexual species and one hybrid taxon with diploid and triplid individuals had been collected during recent research projects across Europe in accordance with environmental protection legislation. The use of frog material was supervised under the permit number CZ 02361 certified and issued by the Ministry of Agriculture of the Czech Republic and followed European standards in agreement with §17 of the Act No. 246/1992 coll., and institutional and national guidelines. For design of a new molecular marker and tests of genome elimination pathways onchromosomal levels we used individuals from four Central-European populations (*P. ridibundus*, *P. lessonae* and *P. esculentus*), one Eastern-European population (triploid *P. esculentus*) and four Southern-European populations (*P. kurtmuelleri*, *P. shqipericus*, *P. epeiroticus*). Other 10 juvenile/subadult *P. esculentus* individuals were F1 descendants of the above-mentioned adult parental taxa, produced via laboratory crosses (**Supplementary Table S1**). A power of studied cytogenetic markers in relation to its species-specifity was tested at several individuals per taxon over their geographic distribution ranges. The artificial fertilization procedure is described in Doležálková-Kaštánková et al., (2022).

Adult frogs were taxonomically pre-determined based on external morphological characters (Günther, 1990; Papežík, 2021) and later verified genetically using previously determined and routinely applied molecular markers involving one Sanger-sequenced mitochondrial DNA marker (*ND2* gene) and 11 nuclear DNA microsatellite markers. These were compared to previously published data to confirm their taxonomical identification following Doležálková-Kaštánková et al., (2021). Total genomic DNA was extracted from an interdigital forelegs membrane using a commercial Tissue DNA Isolation Kit (Geneaid Biotech, Taipei, Taiwan) following a manufacturer’s protocol.

### RepeatExplorer2: Data processing and graph-based clustering

Obtained genomic DNA was used to prepare Illumina paired-end libraries with 450-bp inserts and then sequenced on the Illumina NovaSeq 6000 platform at Novogene (HK) Co., Ltd. (Hong Kong, China). A total of 9.64 Gb of P. lessonae and 8.22 Gb of P. ridibundus raw data were obtained from short-read sequencing, representing 1.53× and 1.10× genome coverage, respectively, given the genome size estimates provided by Vinogradov (1998).

The resulting 150-bp paired-end reads were filtered using Cutadapt (version 1.15; Martin 2011), using the ‘—nextseq-trim’ option for two-color chemistry, using the following parameters: ‘-j0 -bGATCGGAAGAGCACACGTCTGAACTCCAGTCACATCACGATCTCGTATGCCGTCT TCTGCTTG -bAATGATACGGCGACCACCGAGATCTACACTCTTTCCCTACACGACGCTCTTCCGA TCT -BGATCGGAAGAGCACACGTCTGAACTCCAGTCACATCACGATCTCGTATGCCGTCT TCTGCTTG -BAATGATACGGCGACCACCGAGATCTACACTCTTTCCCTACACGACGCTCTTCCGATCT --nextseq-trim=20 --length=130 --minimum-length=130’ and for the second pass (see below): ‘-nextseq-trim=20 -m 150 -a AATGATACGGCGACCACCGAGATCTACACTCTTTCCCTACACGACGCTCTTCCGA TCT-AGATCGGAAGAGCACACGTCTGAACTCCAGTCACNNNNNNATCTCGTATGCCGTCTTCTGCTTG’. A total of 300,000 read pairs (approx. 0.01× genome coverage) for each species were subsampled from the corresponding datasets and interlaced using the RepeatExplorer2 pipeline tools (Novák et al., 2013). Finally, repeat analysis was performed usingthe RepeatExplorer2 pipeline, including the tandem repeat a nalyzer (TAREAN) (Novák et al. 2013, Novák et al. 2020). The algorithm was set to use Metazoa version 3.0 protein domain database and perform comparative analysis, as well as automatic filtering of abundant repeats. Results were processed in R and RStudio (R Core Team 2020, RStudio Team 2019) using an in-house script (for details, see Supplementary File S1).

In order to achieve a better resolution of low-abundance clusters, a second pass of the RepeatExplorer2 analysis was performed with several adjustments. All contigs for the 25 most abundant repeats identified by RepeatExplorer2 in the previous run were extracted, and corresponding reads were removed from the data using Bowtie2 (version 2.3.5.1; Langmead and Salzberg, 2012) with the following setting: ’-p 3 --fast-local -f --un-conc’. Only unmapped reads were kept and converted to fasta with FASTX-Toolkit (version 0.0.14; Hannon, 2010). Three technical replicas for each species were created by subsampling corresponding datasets to approximately 0.03× coverage with Seqtk (version 1.0; Li, 2013). Seed number was generated pseudo-randomly with Python (version 3.4.1; Python Software Foundation 2020) and each replica contained 608,334 read pairs, corresponding to 0.02×–0.03× genome coverage. Finally, datasets were interlaced in RepeatExplorer2 Galaxy-based web server (Novák et al., 2013) and repeats were analyzed using RepeatExplorer2 with the same parameters as previously.

### Primer design

To amplify the repetitive element enriched in *R. ridibundus*, custom primers were designed in Geneious Prime 2020.1.2 (https://www.geneious.com). First, all reads corresponding to the element in the second pass of RepeatExplorer2 were extracted and used for de-novo assembly with Geneious assembler and default settings. The longest contig (1,976 bp) was investigated for the presence of conserved protein domains by DANTE with Metazoa 3.1 protein domain database (Neumann et al. 2019). A pair of primers was designed manually to avoid the identified motifs and to achieve as long amplicon as possible (1,529 bp). Primer sequences are listed in **Table 1**.

**Table 1.**
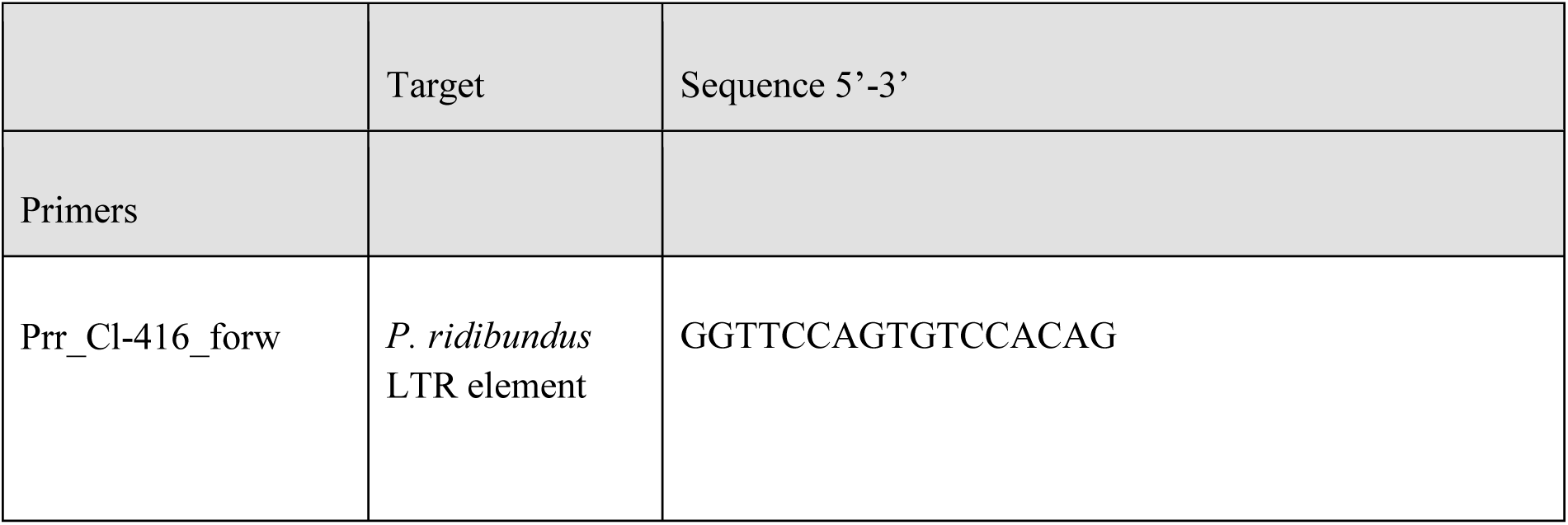
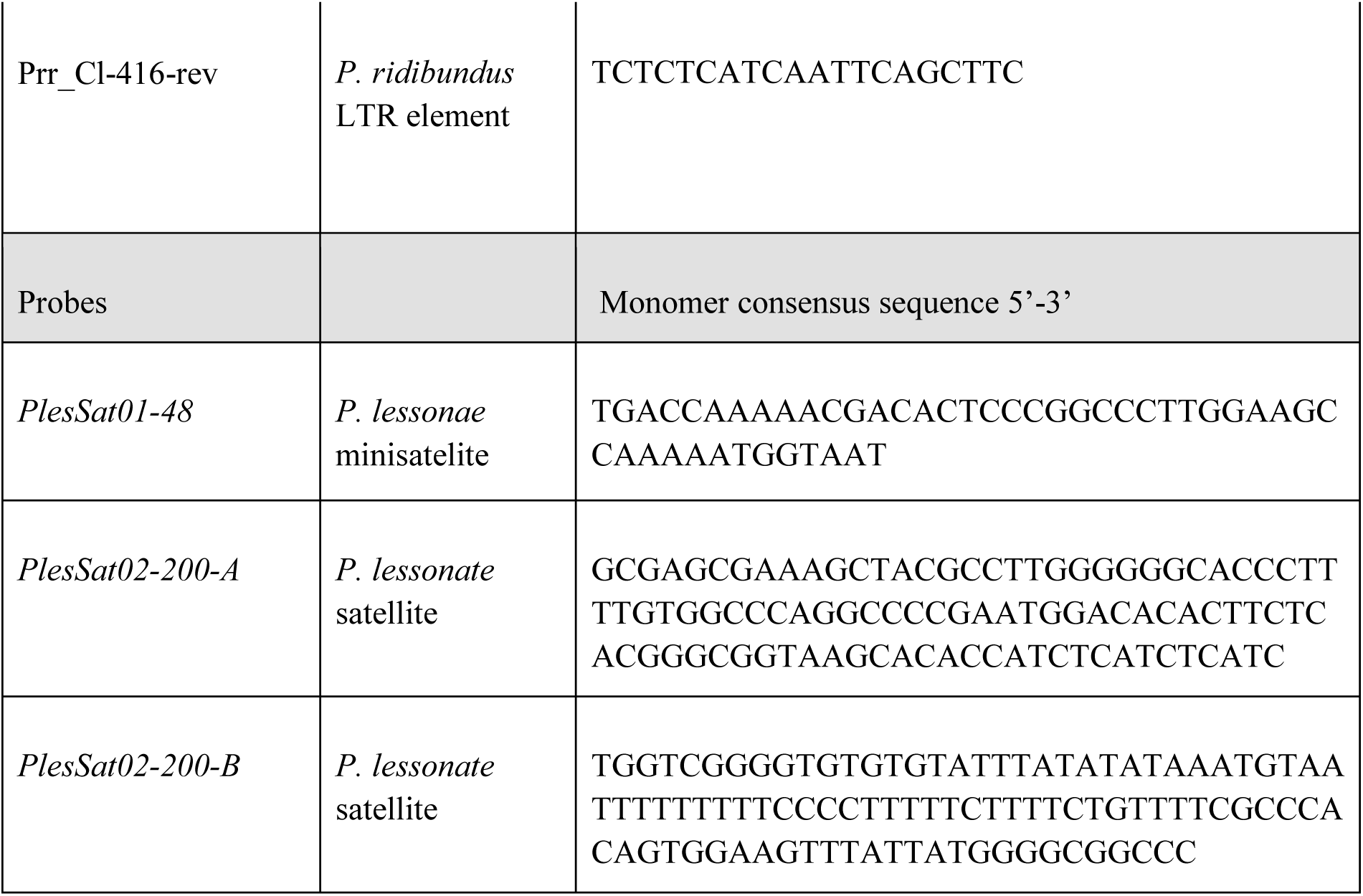
Primers and probes used in this study.

### Probes for fluorescent *in situ* hybridization (FISH)

For detection of the *R. lessonae* enriched satellites, three custom-made oligonucleotides labelled with Cy3 at the 5’ end were synthesized by Generi Biotech (Hradec Králové, Czech Republic). The probes covered the whole monomer length of *PlesSat01-48* minisatellite (48 bp), and in the case of *PlesSat02-200*, two 97-bp-long probes (*PlesSat02-200-A* and *PlesSat02-200-B*) were generated to encompass almost the entire monomer length (which is 200 bp). For the repeat probe sequences, see **Table 1**.

### Chromosome preparation, probe labelling and fluorescence *in situ* hybridization

Chromosome spreads were prepared from peripheral whole blood leucotytes using the protocol outlined by Johnson Pokorná et al., (2016) for reptiles, with the only modification being the cultivation temperature set at 22.5 °C. Labelling of *RrS1* probe specific to the (peri)centromeric tandem repeat of *P. ridibundus* (Ragghianti et al., 1995) and newly developed *PlesSat01-48* was performed by PCR (annealing temperature 62 °C) with the genomic DNA of *P. ridibundus* and *P. lessonae* correspondingly. Primers used for labelling *RrS1* probe were used from Ragghianti et al., (1995): the forward 5’-AAGCCGATTTTAGACAAGATTGC-3’, the reverse 5’-GGCCTTTGGTTACCAAATGC-3’. Primers used for labelling *PlesSat01-48* probe were designed in Geneious Prime 2020.1.2 (https://www.geneious.com): the forward 5’-TTTGGCTTCCAAGGGCCGGG-3’, the reverse 5’-TGACCAAAAACGACACTCCC-3’. Primers used for labelling *PlesSat02-200* probe were: the forward: 5’-GCGAGCGAAAGCTACGCCTTGGG-3’, the reverse: 5’-GCTTACCGCCCGTGAGAAGTG-3’. *RrS1* probe was labelled with biotin-16-dUTP, *PlesSat01-48* and *PlesSat02-200-B* were labelled with digoxigenin-11-dUTP (both Roche, Mannheim, Germany).

FISH was performed according to previously published protocol (Dedukh et al., 2019, 2020). We used two PCR-labelled probes for *RrS1* and *PlesSat01-48* repeats and only PCR labelled probes for *PlesSat02-200* satDNA. The hybridization mixture per slide contained 50% formamide, 10% dextran sulfate, 2× ЅЅС (saline-sodium citrate buffer; 20 × SSC – 3 M NaCl 300 mМ Na_3_C_6_H_5_O_7_), 200 ng of each labelled probe and 10-fold excess of salmon sperm DNA (Sigma-Aldrich) in relation to the amount of the probe. Hybridization mixture was spotted onto slides to perform simultaneous denaturation of the probe and chromosomal DNA which was held at 75 °C for 5 min. Slides were then incubated for 12–24 h at room temperature (RT) in a humid chamber. After hybridization, slides were washed three times in 0.2× SSC at 50 °C for 5 min. The uridines of the probe conjugated with biotin and digoxigenin were detected using streptavidin-Alexa 488 (Invitrogen, San Diego, CA, USA) and anti-digoxigenin-rhodamine (Invitrogen, San Diego, CA, USA), respectively. Afterwards, the slides were dehydrated in an ethanol series (50 %, 70 % and 96 %, 2 min each) and air-dried. The chromosomes were counterstained with Vectashield/DAPI (1.5 mg/mL) (Vector, Burlingame, CA, USA).

### Whole mount FISH

Prior to the whole-mount FISH, gonadal tissues were incubated in 0.5% solution of Triton X-100 in 1× PBS for 4-5 hours at RT and washed in 1× PBS for 15 min. Afterwards, tissues were impregnated with 50% formamide, 10% dextran sulfate, and 2× SSC for 3–4 h at 37°C. Hybridization mixture for both PCR labelled probes contained 50% formamide, 2× SSC and 10% dextran sulfate, 20 ng/µl of probes (10 ng of RrS1 probe labelled with biotin and 10 ng of *PlesSat01-48* probe labelled with digoxigenin) and 10-50-fold excess of salmon sperm DNA. Probe for *RrS1* and *PlesSat01-48* repeats were labelled by PCR. Gonadal tissues were placed in the hybridization mixture, followed by denaturation at 82 °C for 15 min. Tissues were incubated for at least 24 h at RT. Afterwards, gonadal tissues were washed three times in 0.2× SSC at 50 °C for 15 min each and transferred to 4× SSC containing 1% blocking reagent (Roche) for 1 h at RT. Biotin was detected by avidin conjugated with Alexa 488 (Invitrogen, San Diego, CA, USA); digoxigenin was detected by anti-digoxigenin antibodies conjugated with rhodamine (Invitrogen, San Diego, CA, USA). After incubation for 12 h with antibodies against digoxigenin and streptavidin, tissues were washed three times in the 4× SSC for 15 min each. The tissues were stained overnight at RT using DAPI (1 µg/µl) (Sigma) diluted at 1× PBS.

### Confocal laser scanning microscopy

Prior to scanning, tissues were placed in a drop of Vectashield antifade solution. Gonadal fragments were analyzed using Leica TCS SP5 confocal laser scanning microscope based on the inverted microscope Leica DMI 6000 CS (Leica Microsystems, Germany). We used HC PL APO 40× objective for the scanning of whole gonadal tissues. Diode, argon and helium-neon lasers were used to excite the fluorescent dyes DAPI, fluorochromes Alexa 488 and rhodopsin, respectively. The capture of images and their further processing were performed in LAS AF software (Leica Microsystems, Germany). 3D-volume rendering and surface reconstructions of confocal image stacks were performed in Imaris 7.7.1 (Bitplane). The image stacks used for reconstructions were cropped to region of interest (ROI), preserving image voxel dimensions set during image acquisition. Isosurfaces of multi-channel images were created for each channel separately, applying automated thresholding parameters for channel intensity cutoffs. ROI isosurfaces were split into separate surface objects corresponding to individual nuclei (for DAPI channel) or FISH signals (Cy3 channel). In some cases, when two adjacent nuclei were inseparable as individual surface objects, isosurface reconstruction was performed via the “manual creation” tab. Only surface objects belonging to individual germ cells were kept on the reconstruction for highlighted visualisation of germ cells.

## RESULTS

### Marker discovery

The first pass RepeatExplorer2 analysis identified seven highly enriched (|log2| ratio > 4) repeats in *P. lessonae* and five in *P. ridibundus* (not shown). These results were confirmed by the second analysis. Only the most enriched elements for each species were chosen for further study, representing two satellite markers and one retrotransposon marker for *P. lessonae* and *P. ridibundus*, respectively. Detailed information for each repeat of interest is provided in **Table 2**.

**Table 2.** List of markers identified by RepeatExplorer2 for *Pelophylax lessonae* and *P. ridibundus*.

| ID | Species | REx2 annotation |
| --- | --- | --- |
| *PlesSat01-48* | *P. lessonae* | minisatellite, monomer 48 bp |
| *PlesSat02-200* | *P. lessonae* | satellite, monomer 200 bp |
| *Prr_Cl-416* | *P. ridibundus* | BEL/Pao retrotransposon |

### Novel *PlesSat01-48* tandem repeat is specific for *P. lessonae* and closely related species

We mapped two selected repeats identified by RepeatExplorer2 analysis, *PlesSat01-48* and *PlesSat02-200,* on chromosomes of parental species as well as hybrids. The *PlesSat01-48* probe hybridized to (peri)centromeric region of two pairs of *P. lessonae* chromosomes. Specifically, it was observed on the *P. lessonae* acrocentric chromosome pair 8 and the NOR-bearing chromosome pair 10, while absent on *P. ridibundus* chromosomes (**Figure 1** **a, b**). Dual-color FISH revealed *RrS1* repeat to be accumulated specifically in the (peri)centromeric regions of all *P. ridibundus* chromosomes and none *P. lessonae* chromosome, while *PlesSat01-48* repeat occupied exclusively *P. lessonae* chromosomes (**Figure 1** **a, b**).

**Figure 1.**
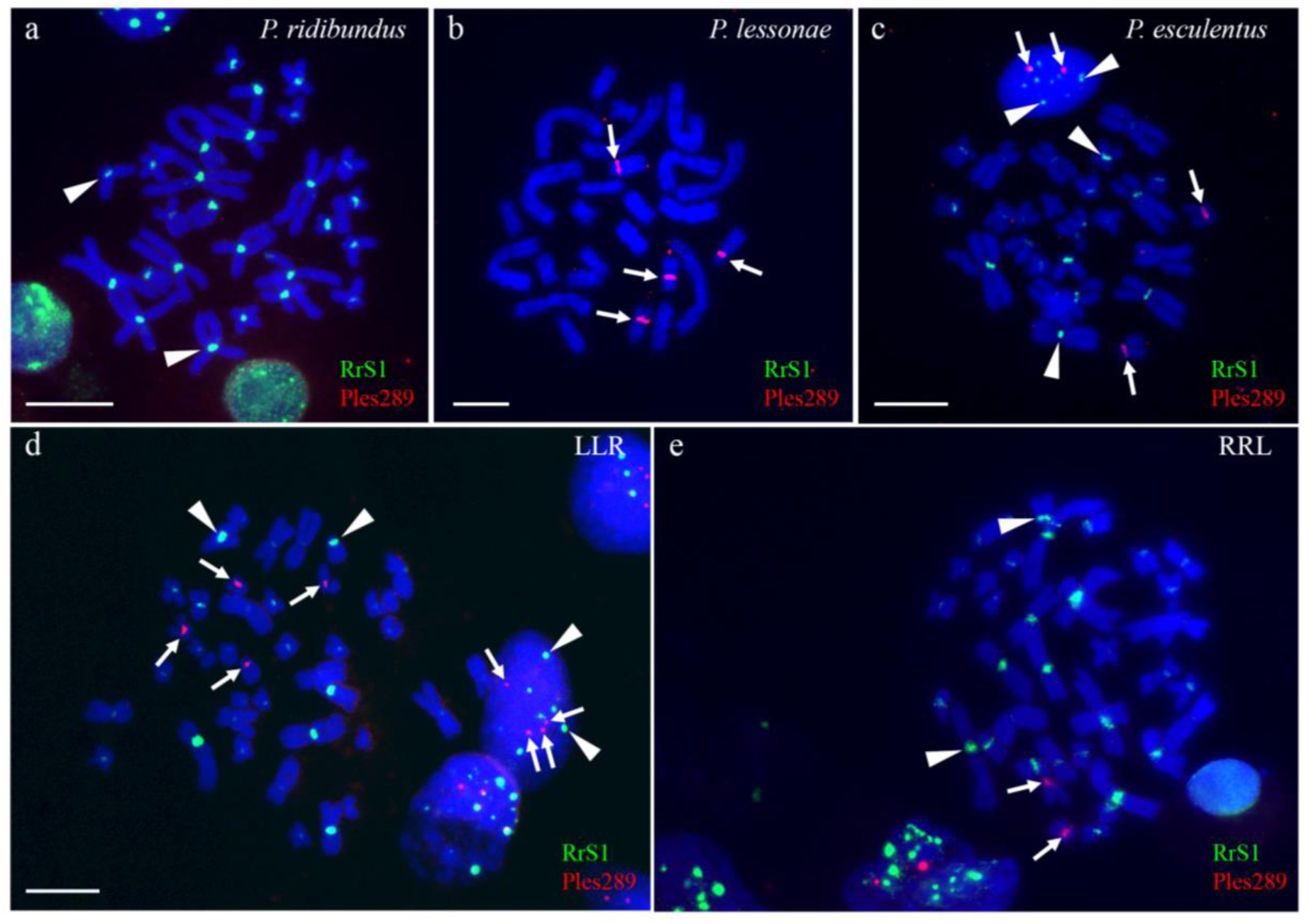
FISH-based mapping of *RrS1* and *PlesSat01-48* tandem repeats on chromosomes of *P. ridibundus* RR (a), *P. lessonae* LL (b), diploid *P. esculentus* (c) and triploid LLR (d) and RRL (e) genotypes. Tandem repeat *RrS1* is specific for (peri)centromeric regions of all chromosomes of *P. ridibundus* (indicated by arrowheads), while novel tandem repeat *PlesSat01-48* (indicated by arrows and coded as Ples289 in the Figure) is localized in (peri)centromeric regions of two pairs of *P. lessonae* chromosomes. In interphase nuclei of diploid **(c)** and triploid **(d)** hybrids *P. lessonae* and *P. ridibundus* chromosomes are clearly identified also in cell nuclei during interphase. Scale bars = 10 µm.

Using FISH with the probes corresponding to the *RrS1* and *PlesSat01-48* (peri)centromeric repeats, we clearly distinguished *P. ridibundus* and *P. lessonae* chromosomes in metaphase plates of diploid as well as triploid hybrids with LLR and RRL genome compositions (**Figure 1** **c - e**). Moreover, such an approach served as a reliable method to identify centromeres of both parental species not only at the metaphase stage but also in interphase nuclei (**Figure 1** **c, e**) and spermatids (Supplementary **Figure S1**).

Mapping of *PlesSat02-200* tandem repeat revealed its localization in the interstitial site of the long arms of chromosome pair 6 or 7 (further denoted as chromosome 6/7) of *P. ridibundus* and *P. lessonae,* as precise identification of the particular chromosomes was difficult (**Supplementary Figure S2 a**). Moreover, in *P. lessonae* we detected this probe also on the short arms of chromosome 6/7 and on the short arms of the chromosome pair 4 (**Supplementary Figure S2 b**). However, the signal on chromosomes 6/7 and 4 was inconsistent and weak. In *P. esculentus*, we detected bright signals on both chromosomes 6/7 and additional weak signals on one chromosome 4 and chromosome 6/7 (**Supplementary Figure S2 c**). Considering that the probe for *PlesSat02-200* detected chromosomes of both parental species and gave inconsistent signals on chromosomes 4 and 7 (or 6), we omitted it for further analyses.

Additionally, we applied FISH with probes for the *RrS1* and *PlesSat01-48* (peri)centromeric repeats to chromosomes of other species from the genus *Pelophylax*, namely, *P. kurtmuelleri*, *P. epeiroticus*, *P. perezi*, *P. shqipericus* and *P. bergeri* (**Figure 2**). We observed the presence of *RrS1* repeat in (peri)centromeric regions of all chromosomes in these species (**Figure 2** **a - e**). The novel tandem repeat *PlesSat01-48* was found only in (peri)centromeric regions of two chromosome pairs in *P. shqipericus* and *P. bergeri* (**Figure 2** **d, e**).

**Figure 2.**
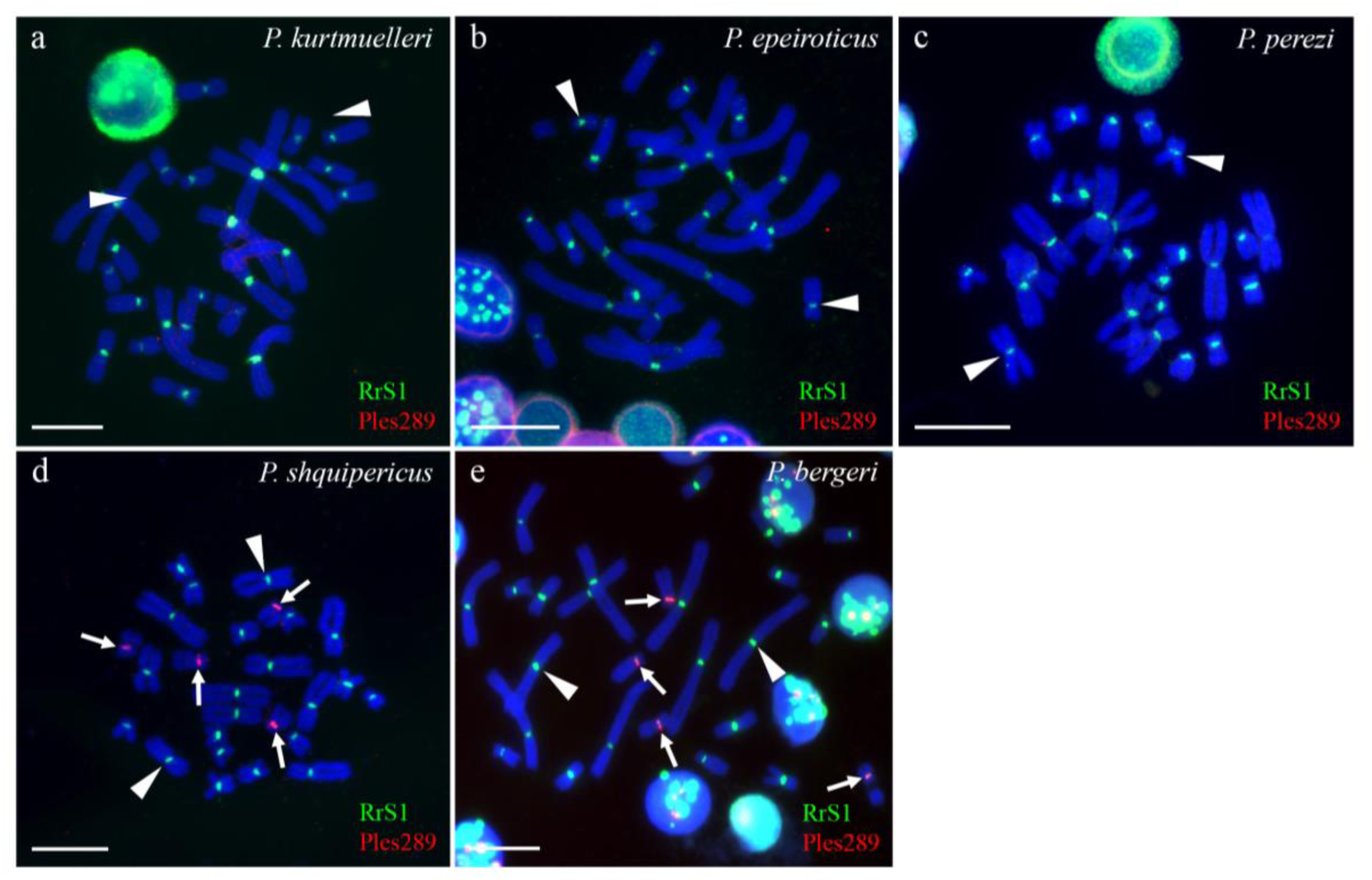
Mapping of *RrS1* and *PlesSat01-48* tandem repeats on chromosomes of *P. kurtmuelleri* (a), *P. epeiroticus* (b), *P. perezi* (c), *P. shqipericus* (d) and *P. bergeri* (e). Tandem repeat *RrS1* (indicated by arrows) is localized in (peri)centromeric regions of all chromosomes in studied species; novel tandem repeat *PlesSat01-48* (coded as Ples289 in the Figure) is localized in (peri)centromeric regions of two pairs of *P. shqipericus* and *P. bergeri* chromosomes. Scale bars = 10 µm.

### In the gonocytes of diploid F1 hybrid tadpoles, micronuclei, misaligned and lagging chromosomes were preferentially presented on *P. lessonae* chromosomes

Dual-color FISH with *RrS1* and *PlesSat01-48* probes were applied to the intact gonadal tissue of 13 F1 hybrid tadpoles (Gosner stages 28–38) obtained from three crosses of *P. ridibundus* female and *P. lessonae* male (crosses ID 21-2020, 37-2020, and 12-2022), two tadpoles from one cross of *P. lessonae* female and *P. ridibundus* male (cross 57-2020) (**Supplementary Table S1**). In addition, we analyzed three tadpoles from one cross of *P. esculentus* female with *P. lessonae* male (cross 27-2020) (**Supplementary Table S1**). We observed germ cells in all tadpoles from crosses of *P. esculentus* female and *P. lessonae* male (cross 27-2020). In the majority of the interphase nuclei of gonocytes, we found approximately 13 signals of *RrS1* probe corresponding to the haploid chromosomal set of *P. ridibundus* (n=13) (Figure 3 a,b). The number of signals generated by the *PlesSat01-48* probe ranged from none to two. These results suggest gradual elimination of *P. lessonae* genome from interphase nuclei of gonocytes. In total, we examined 2 584 gonocytes and found 208 cells with micronuclei. In micronuclei, we detected *P. ridibundus* centromeres in 6,1 % micronuclei (n=15), whereas in the vast majority of micronuclei, we have not reveal any signals (75,6 %, n=186) and detected *P. lessonae*-specific repeat in 18,3 % of micronuclei (n=45). We found four mitotic plates with misaligned chromosomes (7 %) which showed an abnormal attachment to the spindle.

**Figure 3.**
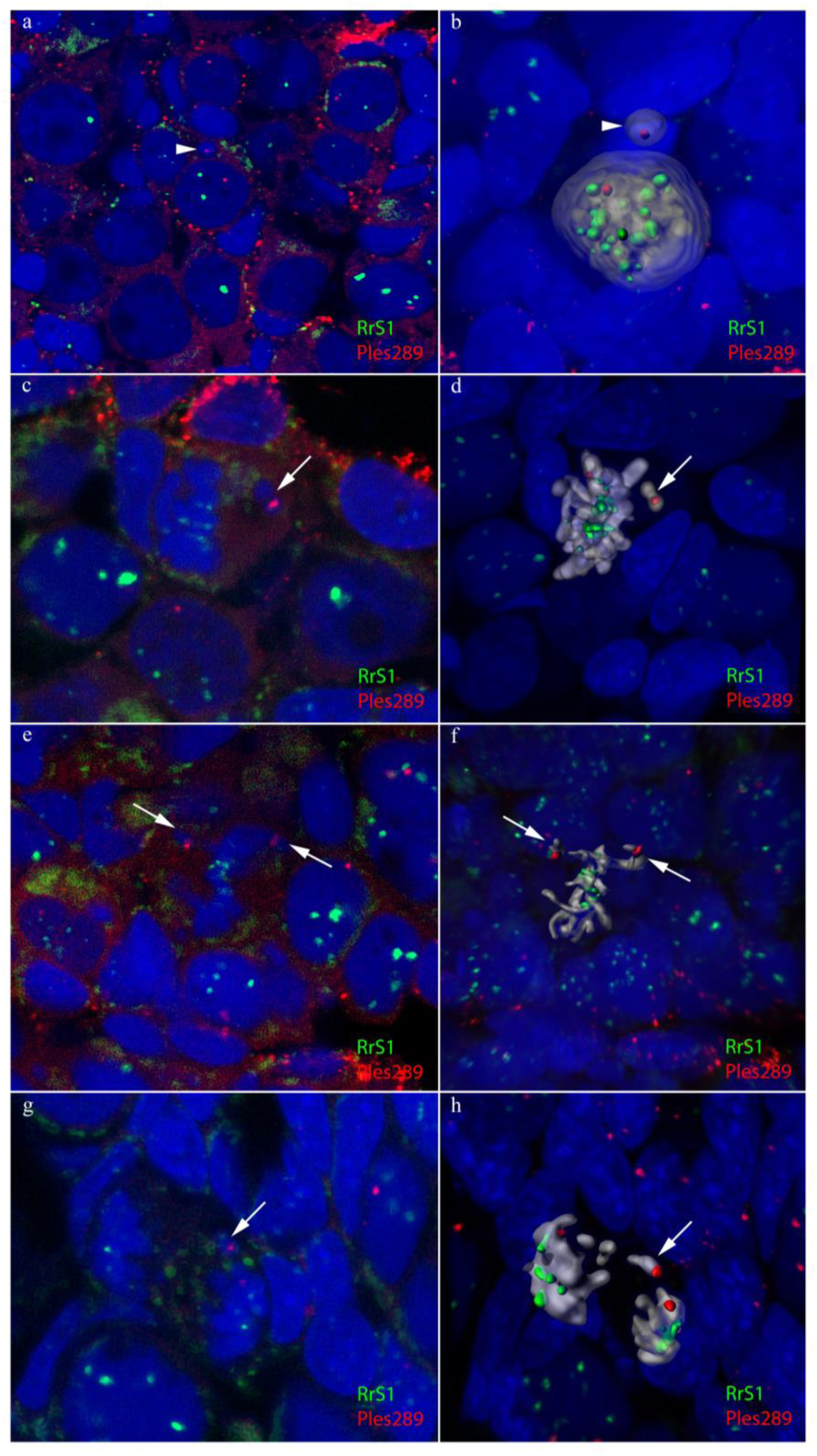
Detection of *P. lessonae* chromosomes in micronuclei (a, b), misaligned chromosomes (c, d, e, f) and lagging chromosomes (g, h) in F1 hybrids. Two-colour FISH with probes specific to *RrS1* and *PlesSat01-48* (coded as Ples289 in the Figure) tandem repeats revealed *P. ridibundus* and *P. lessonae* chromosomes in the cell nuclei and only *P. lessonae* centromere in the micronuclei which is indicated by arrowhead **(a)**. 3D surface reconstruction clearly demonstrates the presence of one *P. lessonae* centromere (red) in the micronucleus (shown by arrowhead) and 13 *P. ridibundus* centromeres (green) in the interphase nucleus (gray) of diploid F1 hybrid **(b)**. Misaligned (c, e) *P. lessonae* chromosomes (indicated by arrows) were observed during the metaphase stage and 3D surface reconstruction of these chromosomes (with red centromere) is presented in images **d, f.** Lagging (g, h) *P. lessonae* chromosome (indicated by arrow) during the anaphase stage was visualised after two-colour FISH with 3D surface reconstruction of lagging chromosome (with red centromere) presented in image **h**. Images **a, c, e, g** are single confocal sections of 0.7 µm in thickness. Scale bars = 10 µm.

Similarly to tadpoles obtained from a cross of *P. esculentus* female with *P. lessonae* male, in all tadpoles from crosses of *P. ridibundus* female and *P. lessonae* male (crosses ##37-2020, 21-2020, 12-2022), we observed standard istribution of gonocytes. Gonocytes usually possessed a haploid *P. ridibundus* genome, and the number of signals from *PlesSat01-48* probe ranged from none to two (Figure 3). Out of 14 362 examined gonocytes we found 1 005 cells with micronuclei in the cytoplasm (Figure 3). In the vast majority of micronuclei, we did not observe signals from any of the applied probes (79,1 %, n=1 523), while the *P. lessonae*-specific repeat was detected in 11,7 % of micronuclei (n=225). Only in 9,2 % (n=177) of micronuclei, we detected *P. ridibundus* centromeres (Figure 3). We found 44 mitotic plates with misaligned chromosomes (20,4 % of the entire analyzed set). Out of the total 60 observed misaligned chromosomes, 13 (21,7 %) belonged to *P. ridibundus* genome, 18 (30 %) were of *P. lessonae* origin, and remaining 29 ones did not carry any hybridization signal (Figure 3). Moreover, we observed eight cells at anaphase stage to possess lagging chromosomes. Out of all lagging chromosomes (n=15), six bore a signal from *P. lessonae*-specific probe, and nine displayed no signal (Figure 3). Misaligned and lagging chromosomes without any signal were considered as *P. lessonae* chromosomes, based on the previous findings by Dedukh et al., (2020) showing that micronuclei without the signal carry typically *P. lessonae* chromosomes or acentric fragments. Thus, tadpoles obtained from crosses of *P. ridibundus* female and *P. lessonae* male frequently eliminated *P. lessonae* chromosomes, which could not attach to the spindle and therefore were lagging during anaphase.

Interestingly, we observed a decreased number of germ cells distributed unevenly across the gonadal tissue in two analyzed tadpoles from one cross of *P. lessonae* female and *P. ridibundus* male (cross 57-2020). The gonads consisted of large clusters formed by somatic cells only. In the interphase nuclei of gonocytes, we detected a variable number of *P. ridibundus* chromosomes which frequently varied from 5 to 13. Furthermore, we observed *PlesSat01-48* probe signals ranging from none to two, where the letter was the most abundant pattern in the analyzed set of cells. After analysing 159 germ cells, we found 20 of them to possess micronuclei. Fifteen out of 26 micronuclei included *P. ridibundus* chromosomes. The rest of them did not exhibit any signal, which suggests that these micronuclei most likely contained *P. lessonae* chromosomes. We did not find misaligned or lagging chromosomes in these tadpoles.

### Phylogenetic distribution of hybridization signals over European *Pelophylax* species generated by *P. ridibundus* and *P. lessonae* species-specific markers

By physical mapping of the *RrS1* and *PlesSat01-48* markers onto metaphase chromosomes of another five European *Pelophylax* species, we obtained a signal distribution pattern over phylogeny of seven taxa using the mitochondrial DNA *ND2* gene tree reconstruction (**Figs. 2a-e** **and** **Figure 4**). The *RrS1* signal appeared on chromosomes from *P. kurtmuelleri*, *P. epeiroticus*, and a basal *P. perezi* (**Figs. 2** **a, b, c**) and also on chromosomes from *P. shqipericus* and *P. bergeri* (**Figs. 2** **d, e**). The novel *PlesSat01-48* repeat was present on chromosomes exclusively at the *Pelophylax* lineage grouping *P. lessonae*, *P. shqipericus*, and *P. bergeri* (**Figs. 2** **d, e and** **Figure 4**).

**Figure 4.**
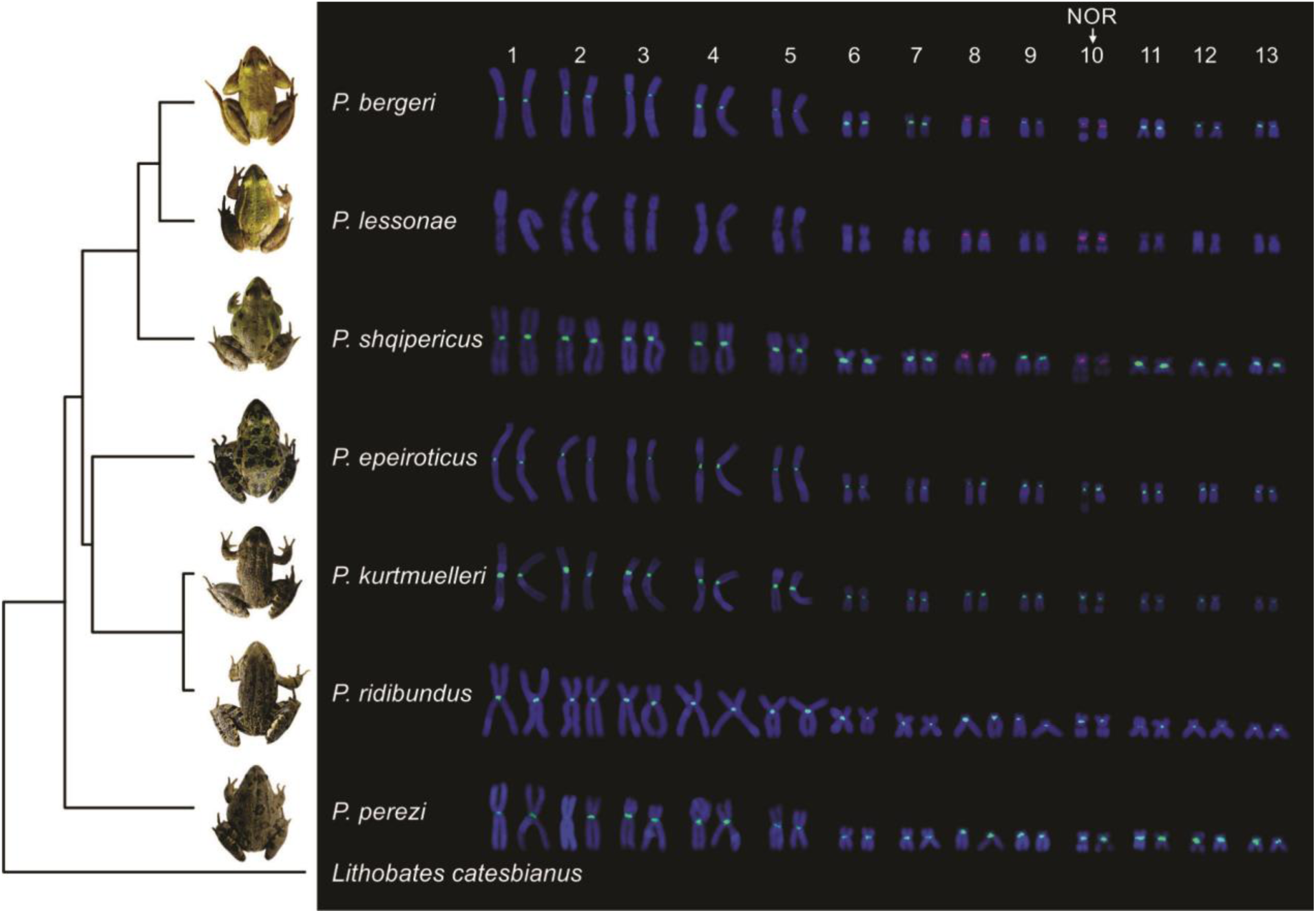
Karyotypes of seven European *Pelophylax* species after two-coloured FISH with *RrS1* and *PlesSat01-48* tandem repeat probes, with distribution of signals across phylogeny using the ND2 gene of the mitochondrial DNA. The *PlesSat01-48* probe hybridized to (peri)centromeric region of two pairs of chromosomes, namely on acrocentric chromosome pair 8 and on the NOR-bearing chromosome pair 10 at *P. bergeri*, *P. lessonae* and *P. shqipericus* (red signal). The *RrS1* signal appeared on all chromosomes from *P. epeiroticus*, *P. kurtmuelleri*, *P. ridibundus*, a basal *P. perezi*, but also on chromosomes from *P. bergeri* and *P. shqipericus* (green signal). A detail information to the phylogenetic tree construction and source sequence data are given in Supplementary File S1.

## DISCUSSION

In the present study, we performed bidirectional crosses (**Supplementary Table S1**) of sexual parental taxa *P. ridibundus* and *P. lessonae* and confirmed hybridogenetic reproduction in the resulting hybrid individuals of *P. esculentus*. This makes *P. esculentus* exceptional among hybridogenetic animals, as, for example, a hybridogenetic fish *Poeciliopsis monacha-lucida* results from unidirectional cross between *P. monacha* females and *P. lucida* males (Quattro et al., 1991; Wetherington et al., 1987). Naturally occurring *P. esculentus* is of polyphyletic origin (Hoffmann, 2015). However, the extent of their ongoing *de novo* formation and the role of the geographic origin of parents are still poorly understood.

Laboratory crossing of parental taxa taken from the current natural populations (this study) and analogous experiment performed on populations collected decades ago (Berger, 1968; Hotz et al., 1985) support the ongoing ability of the genus *Pelophylax* to create a new hybridogenetic *P. esculentus*. Similarly to *P. esculentus*, hybridogenesis in *Poeciliopsis* appears to be induced directly by interactions between hybridizing genomes of parental taxa and has not been restricted to a single historical event, or to an action of specific gene or genome factors as proposed for taxa such as clonal gynogenetic *Poecilia formosa* (Stöck et al., 2010) or parthenogenetic *Aspidoscelis* lizards (Cole et al., 2010; C. C. Moritz et al., 1989). However, previous reconstructions of the origins of *P. esculentus* through crosses between *P. ridibundus* and *P. lessonae* also assumed a strong geographic effect because some hybrids were hybridogenetic, while others exhibited standard meiosis (Hotz et al., 1985). Yet, this conclusion needs to be reconsidered in light of the current distribution of *Pelophylax* taxa in Europe (Dufresnes & Mazepa, 2020; Litvinchuk et al, 2020; Papežík et al., 2023 and our data). The parental *P. ridibundus* which produced non-hybridogenetic hybrids with *P. lessonae* in crosses by Hotz et al., (1985) originated from the Balkans, which means it was, in fact, a sister taxon/lineage *P. kurtmuelleri*.

Laboratory production of *P. esculentus* may be achieved easily, which suggests a probable high transition rates from sexual to asexual reproduction in sympatry. Our laboratory-born F1 hybrids were obtained by crossing parental taxa field-captured from randomly selected populations. That said, we have not observed geographic restrictions in the *de novo* origin of hybridogenetic *P. esculentus* in the context of the Central European region. The closest distance between sites of parental taxa was 35 km, while the largest distance between the collected parental taxa for crosses was 380 km. Temperate Central Europe is known for hybrid zones between eastern and western lineages of various taxa (Jablonski et al., 2021; Nürnberger et al., 2016), which probably reflects the genetic divergence that had accumulated between allopatric populations formed during the ice ages. Because the fine-scale phylogeography of *Pelophylax* taxa in the region is still missing (Hoffmann, 2015; Zeisset & Hoogesteger, 2018), we combined parental individuals from a single or from distinct biogeographic regions, draining into the North Sea and the Baltic Sea. The progeny were viable and morphologically indistinguishable from a wild *P. esculentus* in all families (**Supplementary Table S1**). A similar pattern has been recorded also from regions in which the parental taxa originated from the eastern (Dedukh et al., 2019; A. Svinin et al., 2021) northern (Berger, 1967, 1968; Tunner, 1967, 1973) or western specieś distribution area (Blankenhorn et al., 1971; Hotz et al., 1985). It is, therefore, likely that *P. esculentus* arises *de novo* whenever parental species come into reproductive contact.

Our novel *PlesSat01-48* probe for *P. lessonae* chromosomes combined with *RrS1* probe designed for *P. ridibundus* chromosomes by Ragghianti et al. (1995) showed that synthesized *P. esculentus* progeny is not sexual but hybridogenetic (**Figure 3**) and that *de novo* origin of hemiclonal hybridogenetic reproduction in *P. esculentus* is not restricted in space and time. Another *in situ* escape from sexual to hemiclonal reproduction has been laboratory-verified in the fish *Poeciliopsis* (Schultz, 1973; Vrijenhoek, 1993, 1994; Wetherington et al., 1987) while the only known laboratory synthesis of fully clonal parthenogens from bisexual ancestors was reported in parthenogenetic grasshopper *Warramaba* (White et al., 1977; White & Contreras, 1978) and planthopper *Muellerianella* (Drosopoulos, 1978), and gynogenetic spined loaches of the genus *Cobitis* (Choleva et al., 2012; Marta et al., 2023). Circumstances allowing these hybrid taxa, but not the others, to repeatedly switch from sexual to asexual reproduction (Bialic-Murphy et al., 2020; Cole et al., 2010; Lampert et al., 2007; C. Moritz et al., 1991; Stöck et al., 2010) are yet to be understood.

Molecular data provided indirect evidence on the frequent origin of *P. esculentus* based on a diverse clonal variation of *ridibundus* genomes found in *P. esculentus* (Hoffmann, 2015; Pruvost et al., 2015). Rich clonal diversity is, however, known only from the L-E population system. In contrast, the reverse, i.e. R-E system with all-male hybrids, shares a clonal origin (Doležálková-Kaštánková et al., 2018), suggesting its rare formation in the past time. We were, therefore, interested in whether the synthesized *P. esculentus* progeny eliminates preferentially parental genomes of *P. ridibundus* or *P. lessonae*. In all but one case *P. esculentus* tadpoles eliminated mostly *lessonae* genome and retained *ridibundus* genome in their germ cells, well in agreement with previous studies (Chmielewska et al., 2022; Dedukh et al., 2019; Dedukh & Krasikova, 2022). Neither the direction of the cross nor the origin of maternal mtDNA had any effect on an overrepresentation of *ridibundus* hemiclones. This biased genome elimination requires an occurrence of parental *P. lessonae* as a future donor of *lessonae* genome, meaning that laboratory *P. esculentus*, and likely those newly originated in nature, would typically establish the L-E system, a dominant population type in water frogs over Europe (Hoffmann, 2015; Pruvost et al., 2013; Rybacki & Berger, 2008; A. Svinin et al., 2021; A. O. Svinin et al., 2013), and not the R-E or pure *P. esculentus* populations. Since all *P. esculentus* newborns were diploid and originated from two haploid parental gametes, the L-E system with the occurrence of diploid taxa seems to be evolutionarily ancestral to the L-E system with the occurrence of polyploid *P. esculentus*, and to the pure *P. esculentus* populations that are likely derived from the L-E system with polyploids (Chmielewska et al., 2022; Christiansen, 2005, 2009; Dedukh et al., 2022).

Only *P. esculentus* from the family ID 27-2021 eliminated mostly the *ridibundus* genome, while the *lessonae* genome remained in germ cells. We cannot conclude whether such predispositios would successfully establish a rarely formed R-E system. It is noteworthy, however, that *P. ridibundus* father originated from a source population of the R-E system, in which *P. ridibundus* may have recombinant genotypes with introgressed *lessonae* alleles (Uzzell et al., 1976). If confirmed, a potential role of the *P. ridibundus* male in the induction of a reverse hybridogenetic elimination would bring back into play the possibility that some specific genes determine a type of DNA elimination. Irregular genome elimination carrying *P. ridibundus* chromosome typically occurred in diploid and triploid hybrid tadpoles that emerged from backcrosses between wild-collected hybrids and parental individuals (Dedukh et al., 2020). We additionally noticed that the number of micronuclei with *P. ridibundus* chromosomes varies between cross directions. In tadpoles from crosses of *P. lessonae* female and *P. ridibundus* male, we observed micronuclei mostly with *P. lessonae* genome. However, in tadpoles from crosses of *P. ridibundus* female with *P. lessonae* male, we found micronuclei mostly with *P. ridibundus* chromosomes. Interestingly, in the latter crossing arrangement, the tadpoles possessed a decreased number of germ cells compared to reciprocal crosses. Such irregularities may alternatively explain the maternal/paternal effect on a genome selected for elimination.

Here, we showed that even in the F1 generation, interspecific hybrids instantly modify their gametogenesis to establish asexual reproduction via hybridogenesis. These modifications particularly include premeiotic elimination of one of the parental genomes and clonal propagation of the remaining genome due to endoreduplication followed by standard meiosis (*Pelophylax*, *Hypseleotris*) or ameiosis (*Poeciliopsis*) (Cimino, 1972; Majtánová et al., 2021; Ogielska, 1994; D. J. Schmidt et al., 2011; Tunner & Heppich, 1981). In the so far investigated hybridogenetic species, genome elimination occurs gradually during the gonocyte proliferation in the early embryonic development and it does not occur in adult individuals (Chmielewska et al., 2022; Dedukh et al., 2019, 2020; Majtánová et al., 2021). Genome elimination in diploid and triploid hybrids from nature (Chmielewska et al., 2022; Dedukh et al., 2019, 2020) as well as in laboratory F1 hybrids (Dedukh et al., 2019), this study) occurs gradually and involves micronuclei formation. Based on the analysis of micronuclei, it has been suggested that *P. lessonae* chromosomes were eliminated. Yet, it has never been confirmed (Chmielewska et al., 2022; Dedukh et al., 2019, 2020). Here, using a newly developed marker specific for *P. lessonae* chromosomes, we identified *P. lessonae* chromosomes both in micronuclei and in standard cell nuclei during interphase for the first time. During the elimination, gonocyte genomes were suggested to remain aneuploid until one parental chromosomal set is fully degraded (Dedukh et al., 2019, 2020). We demonstrated the presence of aneuploid gonocytes with various numbers of *P. lessonae* chromosomes during the genome elimination stage in F1 tadpoles. Thus, our results for diploid F1 hybrids correspond with earlier data and indicate the ability of selective elimination of *P. lessonae* genome in F1 progeny.

Two hypotheses have been put forward to explain the mechanism of gradual chromosome elimination and micronuclei formation. Micronuclei can be formed via budding from interphase nuclei or from lagging chromosomes during mitosis of gonocytes. Chromosomal lagging is frequently accompanied by chromosomal misalignment in mitotic metaphase. Misaligned chromosomes fail to attach to the spindle and are distributed apart from the rest of chromosome complement which is arranged at the equatorial part of a cell (Potapova & Gorbsky, 2017; Subrahmanyam & Kasha, 1973; Wang et al., 2017). In anaphase, misaligned chromosomes often lag and are thus not incorporated into the daughter cell nucleus after the completion of the cell division. In studied tadpoles, we observed quite a high number of mitotic plates with misaligned chromosomes, suggesting a potential role of chromosomal misalignments in genome elimination (cf. Potapova & Gorbsky, 2017; Subrahmanyam & Kasha, 1973; Wang et al., 2017). Here we note that observation of misaligned chromosomes has been methodologically effective particularly after 3D surface reconstruction, as with standard “2D” FISH it my be more difficult to distinguish whether chromosomes or their pieces positioned apart from metaphase are a result of selective elimination or a technical artifact.

Our data provide strong indication that *P. lessonae* chromosomes misalign with much higher frequency than those of *P. ridibundus*, and then they lag during the remaining parts of mitotic division of the germ cells in tadpoles from crosses of *P. lessonae* female and *P. ridibundus* male. Moreover, we clearly detected lagging of *P. lessonae* chromosomes during the anaphase stage. Thus, in accordance with previous studies (Chmielewska et al., 2022; Dedukh et al., 2019, 2020), we suggest that chromosomes of one of the parental species (usually *P. lessonae*) can be eliminated through the inability of individual chromosomes to attach to the spindle, causing them to lag during the mitotic divisions of gonial cells. The use of the herein designed *P. lessonae*-specific *PlesSat01-48* probe combined with *RrS1* probe previously designed for *P. ridibundus* chromosomes (Ragghianti et al., 1995) may significantly advance investigations of hybridogenesis and programmed DNA elimination and precisely track the trajectories of both parental genomes in hybrid cells at once. Moreover, not only *P. lessonae* but also *P. bergeri* and *P. shqipericus* genomes can be detected in interphase cells and spermatozoa, opening a way to investigate reproductive pathways in other hybrid systems within *Pelophylax* water frogs.

## ACKNOWLEDGEMENTS

The authors would like to thank Sarka Pelikanova (Institute of Animal Physiology and Genetics CAS, v.v.i.) for her help in the laboratory.

## AUTHOR CONTRIBUTIONS

Conceptualization: L.C., A.S., D.D.

Methodology: L.C., A.S., M.A., A.C.V., P.N., D.D.

Investigation: L.C., M.D.K., V.L., A.S., M.A., K.L., A.C.V., M.D., P.N., E.P., A.F., D.D.

Visualization: M.D.K., V.L., M.A., E.P., A.F., D.D.

Funding acquisition: L.C., D.D.

Project administration: L.C.

Supervision: L.C., D.D.

Writing – original draft: L.C., D.D.

Writing – review & editing: L.C., A.S., A.C.V., P.N., D.D.

## DATA AND MATERIALS AVAILIBILITY

All data are available in the main text and the supplementary materials. Sequence reads of the three designed markers will be made available on the SRA (Sequence Read Archive) NCBI upon publication.

