## Supplementary Figures S1 and S2 for "Formation of hemiclonal reproduction and hybridogenesis in *Pelophylax water* frogs studied with species-specific cytogenomic probes"

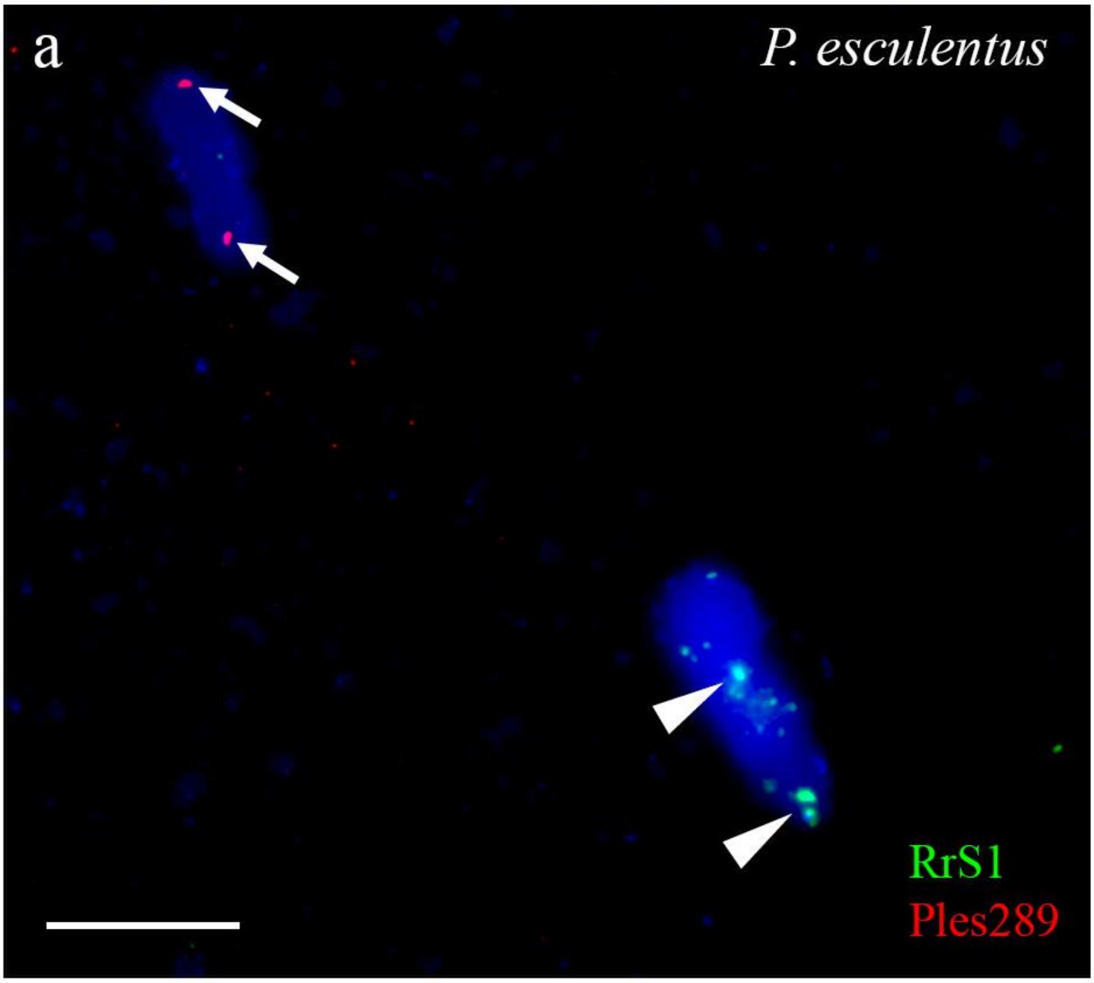


**Supplementary Figure S1. Identification of *P. ridibundnus* and *P. lessonae* genomes in spermatids of diploid hybrid male.** Using a combination of both probes, specific to *P. ridibundus* chromosomes (*RrS1*, indicated by arrowheads) and *P. lessonae* chromosomes (*PlesSat01-48*, coded as Ples289 in the Figure and indicated by arrows) it is possible to identify spermatids with only *P. ridibundus* genome (bottom right) and only with *P. lessonae* genome (top left) simultaneously produced by diploid hybrid male. Scale bars = 10 µm.

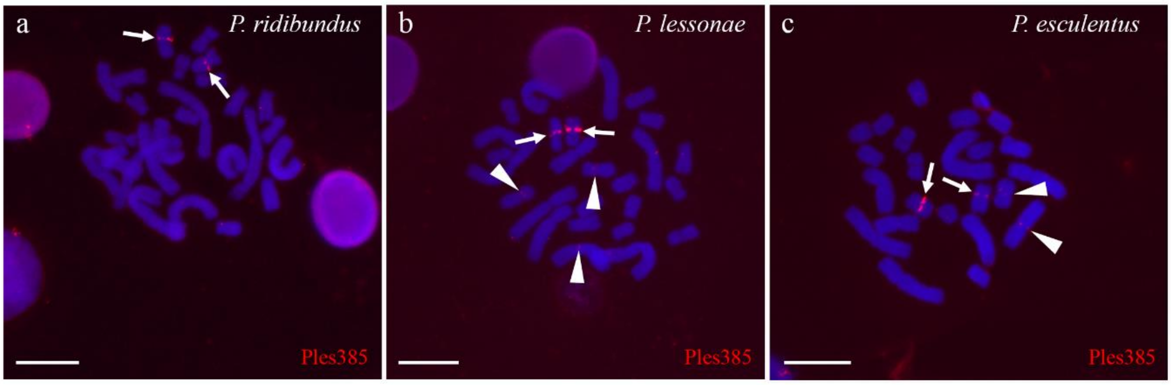


**Supplementary Figure S2. FISH-based mapping of interstitial tandem repeat PlesSat02-200 on chromosomes of *P. ridibundus*, *P. lessonae* and diploid *P. esculentus*.** Tandem repeat *PlesSat02-200* (coded as Ples385 in the Figure) localised in interstitial site of long arms on chromosome 6/7 (see the main text) of *P. ridibundus* (a), *P. lessonae* (b), and *P. esculentus* (c), (indicated by arrows). In *P. lessonae*, it also occupies short arms of chromosome 6/7 and additionally short arms of chromosome 4 (indicated by arrowheads). In *P. esculentus*, we detected both bright and weak signals on homologs of chromosome 6/7 and weak signals on one homolog of chromosome pair 4. Scale bars = 10 µm.
