## Supplementary File S1 for "Formation of hemiclonal reproduction and hybridogenesis in *Pelophylax water* frogs studied with species-specific cytogenomic probes"

**a)** Source sequences of the *ND2* gene (892bp, mitochondril DNA) used fot the phylogenetic tree construction of seven water frog taxa of the genus *Pelophylax*. Codes: bergeri, *P. bergeri* (GenBank no. MN864888), lessonae, *P. lessonae* (original sequence); shqipericus, *P. shqipericus* (original sequence); epeiroticus, *P. epeiroticus* (GenBank no. GU812139); kurtmuelleri, *P. kurtmuelleri* (original sequence); ridibundus, *P. ridibundus* (original sequence); perezi, *P. perezi* (GenBank no. DQ902259); *Lithobates catesbeianus* (Genbank no. EF122830) was used an an outgroup. **b)** Original phylogenetic tree based on the *ND2* mitochondrial DNA of seven European *Pelophylax* species according to Maximul Likelihood analysis, used in the Figure 4. Reference sequences are named as in **a)**.

**a)**

>bergeri

AAAATTCCCCACCCACGAGCTATTGAAGCCGCTACAAAATATTTCCTCACACAAGCTGCT

GCTTCCGCCTTGGTACTATTTTCTAGCCTCATTAACGCCTGACAAACTGGAGAGTGAAAT

ATTGACTCCATCTCTCACTTACCTATAAACGCCCTCTCCATCGCCCTTATGATAAAACTA

GGACTTGCCCCCCTACACTTCTGAATACCTGAAGTCTTACAAGGCATCTCCCTGTCTACA

GGATTAATTTTATCTACCTGACAAAAAATTGCTCCCATAACACTTCTTCTTCAAACCTCT

CACCTAATTAACTTAGATCTGACAATCATCTTAGGCCTCACCTCTATTCTTGTGGGGGGC

TGGGGAGGAATTGGTCAAACTCAATTACGTAAAATTATAGCCTTTTCCTCAATTGGCCAC

CTAGGCTGAATGATTGTTGTTCTAAAATTCAACCCAAGCCTGACCCTCTTTAATTTTATC

TTTTATGTAATCATAACGGCCTCTATATTTTTTTCCTTGATAGCCATTTCCGCCACAAAA

ATATCAGAAATCTCCACCTCCTGATCAAAAACCCCCACTCTCACTACCTCCACATTACTT

ATTCTTCTTTCCCTTGCAGGTCTTCCTCCACTAACAGGCTTTGCGCCTAAATTACTTATT

TCCCTTGAACTGGTAAAACAAAATGCCACTCTCCTTGCCGCCCTAATTATATTGGCCTCC

CTACTAGCCCTATTTTTTTACCTCCGACTTACATATATTGTTTCTCTAACCCTTCCACCC

AATTCCCCTAACTCACTTTCATTCTGACGAACATCAACCAAAACCCCCCCAATTACCGCG

ATTGCAAACACTCTTGCTCTAATTTTTCTCCCAATTACCCCAACCCTCCTTA

>lessonae

AAAATTCCCCACCCACGAGCTATTGAAGCCGCTACAAAATATTTCCTCACACAAGCTGCT

GCTTCCGCCTTGCTACTATTTTCCAGCCTCATTAATGCCTGACAAACTGGAGAATGAAAT

ATTGATTCCATCTCTCATTTACCAATAAACGCCCTCTCCATCGCCCTTATAATAAAACTA

GGACTTGCCCCCCTACACTTCTGAATACCTGAAGTCTTACAAGGCATCTCCCTGTCCACA

GGATTAATTTTATCTACCTGACAAAAAATTGCTCCTATAACACTTCTTCTTCAAACCTCT

CACCTAATCAACTTAGACCTAACAATTATCTTAGGCCTCACCTCTATCCTTGTGGGGGGC

TGAGGAGGAATTGGTCAAACTCAACTACGTAAAATTATAGCCTTTTCCTCGATTGGCCAC

CTGGGCTGAATGATCGTTGTTCTAAAATTCAACCCGAGCCTGACCCTCTTTAATTTTATT

TTTTATGTAATCATAACAGCCTCTATATTTTTTTCCTTAATAACCATTTCCGCCACAAAA

ATATCAGAAATCTCTGCCTCCTGATCAAAAACCCCCACTCTCACTACCTCTACACTACTT

ATTCTTCTTTCCCTAGCAGGTCTTCCGCCACTAACAGGCTTTGCGCCTAAATTACTCATC

TCCCTTGAACTAGTAAAACAAAATGCCACTCTCCTTGCCGCCCTGATTATATTGGCCTCC

CTACTAGCCCTATTTTTTTATCTCCGACTTACATATGTTGTTTCTCTAACCCTTCCACCC

AATTCCCCCAACTCACTCTCATCTTGACGAACATCAACCAAAACCCTACCAATTACCGCG

ATTGTAAACACTCTTGCTCTAATTCTTCTCCCGATTACCCCAACCCTCCTTA

>shqipericus

AAAATCCCCCACCCACGAGCTATTGAAGCCGCTACAAAATACTTCCTTACACAAGCTGCT

GCTTCCGCCTTAGTATTATTTTCTGGCCTCATCAATGCCTGACAAACTGGAGAATGAAAC

ATTGACTCCCTCTCTCATTTACCAATAAATGCCCTCTCCATCGCCCTTATAATAAAACTC

GGACTTGCCCCCCTACACTTCTGAATACCTGAAGTCTTACAGGGCATCTCCCTGTCTACA

GGACTAATTTTGTCTACCTGGCAAAAAATTGCTCCCATGACACTTCTTCTCCAAACCTCT

CACCTAATTAACTTAGACCTAACAATCATCTTAGGCCTCACCTCTATTCTTGTAGGGGGC

TGGGGGGGAATTGGCCAAACTCAACTACGTAAAATTATAGCCTTTTCCTCAATCGGCCAC

CTAGGCTGAATAATTATTATTCTAAAATTTAACCCAAGCCTAACCCTCTTTAATTTTATT

TTTTATGTAATCATAACAGCCTCTATATTTTTTTCCCTAATAGCCATTTCCGCCACAAAA

ATATCGGAAATCTCCACCTCCTGATCAAAAACTCCGACTCTCACTGCCTCCACATTACTT

ATTCTTTTATCCTTAGCAGGTCTCCCACCACTAACAGGCTTTGCACCTAAACTACTTATT

ACCCTCGAACTCGTAAAACAAAGTGCCACTCTCCTTGCTGCCCTAGTTATATTAGCCTCC

CTACTAGCCCTATTTTTTTACCTCCGACTTACGTATGTTATTTCCCTAACCCTTCCACCC

AATTCCCCCAACTCATTTTCGTCTTGACGGACATCAACCAAAACCCCCTCAATTACCGCG

GTTACAAACACTCTTGCCCTAATTTTTCTCCCGATTACTCCAACCCTCCTTA

>epeiroticus

AAAATTCCTCACCCACGAGCCATCGAAGCTGCTACAAAATATTTTCTTACGCAAGCCGCT

GCTTCCGCCTTAGTCCTATTTTCCAGCCTTATTAATGCATGACAAACCGGAGAATGAGGT

GTTGATTCTATCTCCCACCTACCTATAAATGCCCTCTCTATTGCTCTTATAATAAAACTC

GGACTTGCCCCCTTACACTTCTGAATACCTGAAGTCTTACAAGGCATCTCACTACCCACA

GGATTAATTTTATCAACCTGACAAAAAATCGCCCCAATAACGCTTCTTTTACAAACTTCT

CATCTAGCTAATTTACACCTAACAATCATTTTGGGTCTTACTTCTATTCTTATCGGGGGA

TGAGGTGGAATCGGCCAAACTCAACTACGTAAAATTATGGCATTCTCCTCAATCGGCCAT

CTAGGGTGAATAATTGTCATCTTAAAGTTCAATCCGGACTTAACCCTATTTAACTTTGTT

TTTTATATCATCATAACGGCTTCTATATTTTTATCCCTAACTACCATTTCCGCAACAAAA

ATGTCGGAAGTCTCCACCTCATGACCAAAAACCCCCGCCCTTACTGCCTCTACACTACTT

ATTCTCCTATCCTTGGCAGGACTCCCCCCACTCACAGGTTTTGCCCCCAAACTACTTATT

ACCCTCGAACTAGTAAAACAGGACGCCACCCTACTTGCTGCTCTAATTATACTAGCCTCT

CTGCTAGCCCTCTTCTTTTACCTTCGACTAACGTATGTTGTTTCCCTAACTCTCCCGCCT

AACACCCCCAACTCATTTTCCTCTTGACGAACATCAAGTAAATCACTCACAATTACCGCA

ATCATGAATACTCTCGCGCTTGTTCTGCTCCCGCTCACCCCAACCCTCCTCA

>kurtmuelleri

AAAATCCCCCACCCACGAGCCATCGAAGCTGCTACAAAATATTTCCTCACACAAGCTGCT

GCTTCCGCCTTAGTTTTATTTTCCAGCCTCATTAATGCTTGACAAACCGGAGAATGAAAC

ATTGACTCTGTTTCTGACCTACCAATGAATGCCCTCTCCATTGCTCTTATAATAAAACTT

GGACTTGCCCCTCTACACTTCTGAATGCCCGAAGTCTTACAAGGCATCTCACTGTCTACA

GGACTAATCTTATCAACCTGACAAAAAATCGCCCCAATAACACTTCTTCTCCAAACTTCT

CATTTAATTAATTTAGACCTAACAATTATTCTAGGCCTTACCTCTATTCTTGTAGGGGGG

TGAGGGGGGATTGGTCAAACTCAACTACGCAAAATTATAGCATTTTCCTCAATTGGCCAC

CTAGGGTGAATAATTATTGTTCTAAAATTTAATCCAAGCCTGACCCTATTTAACTTTATT

CTTTATATTGTCATAACAGCTTCTATATTTTTATCCCTAATAACCCTTACCACCACAAAA

ATGTCAGAAATTTCTACCTCATGATCAAAAACTCCCACCCTCACTGCCTCTACACTACTC

ATTCTCCTGTCACTAGCAGGACTCCCACCATTAACAGGTTTTGCCCCCAAATTACTTATT

ACCCTAGAACTAGTAAAACAAGATGCTACCCTTCTTGCTGCCCTAATTATATTAGCCTCT

TTACTAGCCCTATTCTTTTATCTCCGACTAACATATATCGTCTCCCTAACTCTCCCGCCA

AACACCCCCAATTCATTTTCCTTTTGACGAACATCAAACAAATCCCTCCCAATTACCGCG

ATTATAAATACTCTAGCACTTATCTTTTTACCACTTACCACAACCCTCCTCA

>ridibundus

AAAATCCCCCACCCACGAGCCATCGAAGCTGCTACAAAATATTTCCTCACACAAGCTGCT

GCTTCCGCCTTAGTTTTATTTTCCAGCCTCATTAATGCTTGACAAACCGGAGAATGAAAC

ATTGACTCTGTTTCTGATCTACCAATGAATGCCCTCTCCATTGCTCTTATAATAAAACTT

GGACTTGCCCCTCTACACTTCTGAATACCCGAAGTCTTACAAGGCATCTCACTGTCTACA

GGACTAATCTTATCAACCTGACAAAAAATCGCCCCAATAACACTTCTTCTCCAAACTTCT

CATTTAATTAATTTAGACCTAACAATTATTCTAGGCCTCACCTCTATTCTTGTAGGGGGG

TGAGGGGCGATTGGTCAAACTCAACTACGCAAAATTATAGCATTTTCCTCAATTGGCCAC

CTAGGGTGAATAATTATTGTTCTGAAATTTAATCCGAGCCTGACCCTATTTAACTTTATT

CTTTATATCGTCATAACAGCTTCTATATTTTTATCCTTAATAACCCTTACCACCACAAAA

ATGTCAGAAATTTCTACCTCATGATCAAAAACTCCCACCCTCACTGCCTCTACACTACTC

ATTCTCCTGTCGCTAGCAGGACTCCCGCCATTAACAGGTTTTGCCCCCAAATTACTTATT

ACCCTAGAACTAGTAAAACAAGATGCTACCCTTCTTGCTGCCCTAATTATATTAGCCTCT

TTACTAGCCCTATTCTTTTATCTCCGACTAACATATATTGTCTCCCTAACTCTCCCGCCA

AACACCCCCAATTCATTTTCCTTTTGACGAACATCAAACAAATCCCTCCCAGTCACCGCG

ATTATAAATACTCTAGCACTTATCTGTTTACCACTTACCAnAACCCTCCTCA

>perezi

AAAACTCCCCACCCCCGAGCCATCGAAGCCGCTACAAAATATTTCCTTACACAAGCTGCT

GCCTCCGCCCTACTTTTATTTTCTAGTCTCATTAATGCTTGGCAAACTGGAGAATGGGCC

ATTAACTCCCTTACTGACCTCCCAATGAATGCTCTCTCCATTGCCCTCATAATAAAACTA

GGACTCGCCCCCCTACACTTTTGAATGCCTGAAGTCCTGCAAGGTATTTCACTATCTACA

GGACTTATTCTATCAACCTGACAAAAAATTGCCCCAATGACACTCCTCCTTCAAACTTCC

CACTTAATTAATCTAAATTTAACAGTTATTCTAGGCCTCACCTCCATTCTTGTAGGCGGG

TGAGGCGGAATAGGTCAAACTCAACTACGTAAAATCATAGCTTTTTCTTCAATCGGCCAC

CTCGGATGAATAATTGTTGTCCTAAAATTTAGCCCAACCCTTACCCTCTTCAACTTTATC

CTCTACATTCTTATAACAGCCTCTATATTTTTATCCCTAATAACCATTTCTGCCACAAAA

ATATCGGAAATCTCCACCTCGTGACCAAAGACCCCCGCCCTCACCGCCACCACACTACTT

GTTCTCCTCTCCCTCGCAGGCCTTCCCCCCCTAACGGGCTTTGCCCCAAAACTGTTAATT

ACCCTAGAGCTAGTAAAGCAAAACGCCACTCTCCTTGCCACCCTAATTATACTAGCCTCC

CTACTAGCCCTATTCTTTTATCTCCGACTGACATACGTCGTCTCCCTAACCCTCCCGCCT

AATACCCCCAACTCATTTTCCTCTTGGCGAACCTCAACCAAACCCCACCCAATAACCGCT

GTCACAAACACCTTTACACTTATCTTCCTCCCCCTTACCCCAACACTCCTCA

>Lithobates catesbeianus

------------------------------------------------------------

------------------ATTTTCAGCCTCATTAGTGCATGACAAACTGGAGAATGAAGT

ATCAACTCTCTTATAGATTTACCAATAAACATCCTCTCTATCGCCCTGATAATAAAACTC

GGTTTGGCCCCGCTACATTTCTGAATGCCTGAAGTTCTTCAAGGAATTTCCCTCCCCACG

GGACTCATCTTATCGACCTGACAAAAAATTGCCCCTATAGCCCTTCTCCTTCAAACTTCT

CACCTTATTAACCTCAACCTAACTATTGCCCTAGGTCTTACATCTATTATAGTTGGAGGC

TGAGGCGGAATTGGTCAAACTCAGCTTCGAAAGATTATGGCCTTTTCTTCTATTGGTCAC

CTAGGATGAATTATTGTCATTTTAAAATTTGATCCACAACTTTCCTTATTAAACTTCGTA

TTATACATTATTATGACAGCCGCCATATTTATATCCCTAACTACCATTTCTGCCACAAAA

ATGTTGGAAATCTCTACCTCATGATCAAAAACCCCTGCTCTCACCACAACCACCATACTT

ATTCTATTATCCTTGGCAGGACTCCCCCCCCTTACAGGCTTTGCCCCAAAACTCTTAATT

ACCCTAGAACTAGTGAAACAAAATGCAACCCTCCTTGCCGTGATAGTCATGTTTATTTCC

CTATTAGCTTTATTTTTTATATCCGACTGACATATGTGGTTACCCTAACTCTCTCCCCA

AATACCCCCAACTCATTACTCACTTGACGAACTGCCTCTCGATCTTACTCAACAACTGCT

ATTATAAATACCATGGCACTTATCCTTCTTCCCATTACCCCAACTCTCCTTC

**b)**

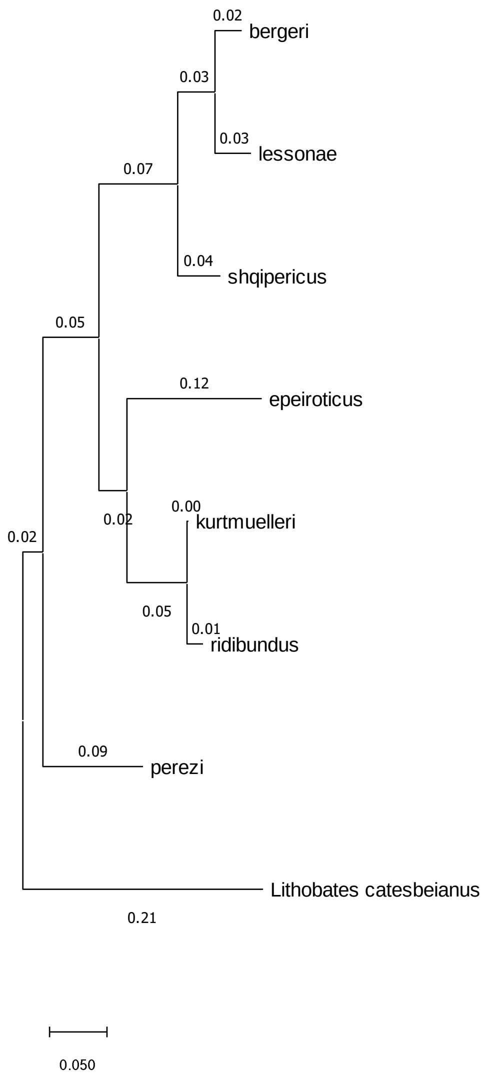
0.02
bergeri
0.03
0.03
lessonae
0.07
0.04
shqipericus
0.05
0.12
epeiroticus
0.00
0.02
kurtmuelleri
0.02
0.05
0.01
ridibundus
0.09
perezi
Lithobates catesbeianus
0.21
0.050
