## Supplementary Table S1 for "Formation of hemiclonal reproduction and hybridogenesis in *Pelophylax water* frogs studied with species-specific cytogenomic probes"

**Supplementary Table S1**: Overview of *Pelophylax* water frog families from artificial crossing experiments.

| | | | Mother´s | | Father´s | |
|---|---|---|---|---|---|---|
| **Frog ID** | **Genotype** | **Sex** | **Genotype** | **Origin** | **Genotype** | **Origin** |
| 21-2020 AD1 | RL | ? | RR | R-R pop | LL | L-E pop |
| 21-2020 AD10 | RL | ? | RR | R-R pop | LL | L-E pop |
| 37-2020 AD9 | RL | ? | RR | R-R pop | LL | L-E pop |
| 37-2020 AD10 | RL | ? | RR | R-R pop | LL | L-E pop |
| 57-2020 AD2 | RL | ? | RL | L-E pop | LL | L-E pop |
| 57-2020 AD3 | RL | ? | RL | L-E pop | LL | L-E pop |
| 27-2021 AD1 | RL | ? | LL | L-E pop | RR | R-E pop |
| 27-2021 AD2 | RL | ? | LL | L-E pop | RR | R-E pop |
| 16-2016 AD5 | RL | M | RR | R-R pop | RL | R-E pop |
| 12-2016 AD5 | RL | M | RR | R-R pop | RL | R-E pop |
